# Interdisciplinary Study on Drug-Induced-Phospholipidosis of Repurposing Libraries through Machine Learning and Experimental Evaluation in Different Cell Lines

**DOI:** 10.1101/2025.07.08.663638

**Authors:** Maria Kuzikov, Adelinn Kalman, Jeanette Reinshagen, Johanna Huchting, Kun Qian, Hanna Axelsson, Marianna Tampere, Päivi Östling, Brinton Seashore-Ludlow, Yojana Gadiya, Philip Gribbon, Andrea Zaliani

## Abstract

**Bigger Picture:** Phospholipidosis is a cellular condition characterized by the excessive accumulation of phospholipids within cells, that also can be induced by medications—especially those known as cationic amphiphilic drugs. While this phenomenon is not always harmful in itself, it can interfere with how cells respond in laboratory experiments and complicate the interpretation of drug screening results, leading to potential delays or failures in the development of new therapies. As the pharmaceutical industry increasingly relies on high-throughput screening and machine learning to accelerate early-stage drug discovery, there is growing interest in identifying compounds that may cause phospholipidosis before they move too far through the development pipeline.

In response to this challenge, we at the EU-Horizon HLTH Remedi4ALL project compiled and analyzed one of the most comprehensive datasets to date—over 5,000 repurposed drugs tested across multiple cell lines—to better understand which compounds are likely to induce phospholipidosis. Using this dataset, we developed a machine learning model capable of predicting the risk of phospholipidosis based on a drug’s chemical structure. By validating our approach across diverse experimental settings, our goal is to provide the broader scientific and pharmaceutical communities with a practical tool to flag problematic compounds early, ultimately helping bring safer, more effective treatments to patients faster.

**Highlights:**

– A comprehensive dataset for a repurposing collection, comprising 5,228 compounds, for induction of phospholipidosis in A549-ACE2 and Vero-E6 cells
–A machine learning model developed using in-house experimental phospholipidosis data across multiple cell lines
–Exploration of chemical features that induce phospholipidosis and investigation of the relationships underlying different cell line susceptibilities
–A case study: correlation of drug-induced phospholipidosis with antiviral compound effects against SARS-CoV-2 using phenotypic screening data

Phospholipidosis (PLD), a cellular adverse effect that is, among others, caused by numerous cationic amphiphilic drugs. Interest is raised within pharma discovery to predict this phenomenon, as it can impact the outcome of phenotypic cellular screens and significantly delay drug development processes. The development of accurate and validated machine learning models for predicting drug-induced PLD across different cell lines and research centers could provide a valuable early application tool for the pharmaceutical industry, potentially accelerating drug discovery and reducing the risk of late-stage failures. We report here the assembly, curation, testing and modeling of one of the largest datasets of repurposed drugs (5K+) tested for PLD induction on different cell lines. A machine-learning classification method was developed and validated to predict whether molecules are prone to induce PLD effects when applied in cell-based screens.

Graphical abstract

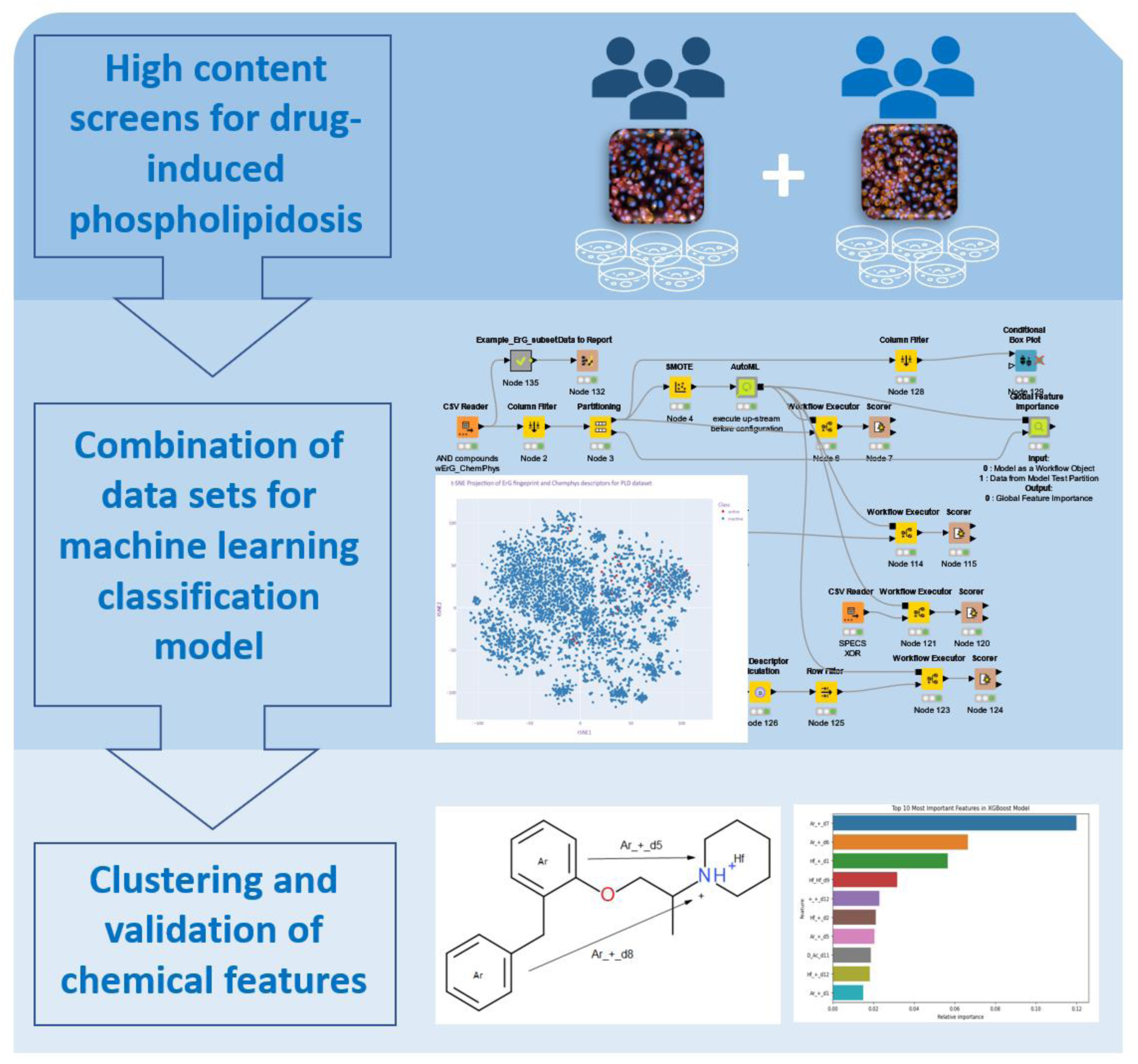

## 1. Introduction

Phospholipidosis (PLD) is a pathologic storage disorder observed as an excessive accumulation of phospholipids in lysosomes^1^. PLD’s involvement in various cellular processes makes it relevant to multiple therapeutic areas, including oncology, neurology, and immunology. The mechanism behind drug-induced PLD is not well understood, it is thought to involve the inhibition of the lysosomal phospholipase activity, an increase of lysosomal pH towards neutral, blockage of mannose-6-phosphate receptor (M6PR) leading to the inhibition of lysosomal hydrolase activities, and changes of lysosomal membrane composition^2,3^.

Special attention is paid when these conditions are induced by chemical entities. Cationic amphiphilic drugs, e.g., chloroquine or tamoxifen, were reported to induce PLD *in vitro* and *in vivo*^4,5^. Emerging evidence suggests that PLD may also impair virus infection and replication cycle *in vitro*^4^. On the one hand, viruses remodel the host’s cellular environment, including phospholipid metabolism, to create optimal conditions for their replication. On the other hand, phospholipids and their regulatory enzymes influence virus replication. For instance, the presence of phospholipids on the plasma membrane prevents the attachment of the respiratory syncytial virus (RSV), while cholesterol oxidation product 25-hydroxycholesterol (25HC) impairs Hepatitis C virus infection by disrupting the formation of the virus-induced double membrane vesicle network necessary for its replication ^6^, ^7^. However, as highlighted by Tunnimo *et al.* for SARS-CoV-2, the antiviral effects associated with drug-induced PLD have not been consistently replicated *in vivo*, leading to concerns that drug-induced PLD may be an artificially generated antiviral effect that may result in the generation of false positive results in cell-based antiviral drug screening^4^. These findings highlight the need for investigation into the chemical properties of compounds that lead to PLD induction in order to minimize false positives that might not have a real protein target as the basis of their action.

Studies on common chemical properties for PLD-inducing compounds have been reported in the literature since 1970s, i.e., the presence of a tertiary amine group that becomes protonated and subsequently trapped inside negatively charged lysosomal vesicles or hydrophobic ring structure with a hydrophilic side chain^8,9,10,11^. Several studies have now demonstrated applications of machine learning (ML) techniques for predicting compounds’ propensity to induce PLD. One of the earliest efforts dates back to 2008, when Ivanciuc trained and compared over 20 different ML models using the Weka software to explore the structure-activity relationship (SAR) behind PLD ^12, 13, 14^. The training set comprised of structural descriptors for 201 compounds, whereof 118 were publicly available and the remaining taken from Pfizer’s internal database. From the comparison, Ivanciuc reported that a Support Vector Machine (SVM) with a Gaussian radial basis function (RBF) kernel achieved the highest performance in a 10-fold cross-validation approach with an accuracy of 97% and Matthews Correlation Coefficient (MCC) of 0.94. Several years later, Lowe *et al.* (2010) expanded the dataset using literature-derived data and developed ML models based on structural and circular fingerprints, as well as a combination of both^15^. They found that circular fingerprints alone led to better models, with Random Forest (RF) and SVM models showing similarly high performance. Their best models achieved over 90% accuracy in predicting PLD-inducing potential. Later, in 2012, Orogo *et al.* constructed a dataset of 743 compounds and investigated the effectiveness of commercially available QSAR software programs in PLD prediction^16^. Their commercial model, which integrated multiple ML approaches, achieved 86% accuracy in predicting PLD. In another study, Fusani *et al.* (2017) combined high-content screening data with *in silico* modeling to predict PLD risk^17^. Using a mix of public and proprietary data from AstraZeneca, they found that the SVM model outperformed RF, achieving 86% accuracy on an external test set. In recent years, interest from both pharmaceutical companies and academia has driven further advances. Boehringer Ingelheim used their *in-vitro* assay results to explore experimental and computational approaches for PLD-inducing compounds^18^, while academic researchers applied cell imaging techniques ^19^. Among these efforts, the AMALPHI (https://www.ba.ic.cnr.it/softwareic/amalphi/) ML platform developed by Lomuscio *et al.* (2024) stands out for its accessibility, as it openly provides the model for broader use^20^. Their best model achieved 88% accuracy and an MCC of 0.76 on the test set, expanding the utility of PLD prediction models beyond traditional user groups. Collectively, these studies have demonstrated two key insights: (i) understanding the relationship between compound structure and PLD can offer valuable clues about PLD-related toxicity, and (ii) larger, more comprehensive datasets enable the discovery of new findings and structural characteristics for compounds PLD effect. As datasets continue to grow and ML techniques evolve, the ability to predict and mitigate PLD-related toxicity will likely improve, leading to safer drug development and more informed therapeutic strategies.

Our findings delve deeper into the risk of false positives associated with PLD in screening, particularly in the context of antiviral drug discovery, and with implications seen during the recent SARS-CoV-2 pandemic. Finally, we make available the ML model, its source code, and training data in public repositories, ensuring openness and data sustainability for future research.

## 2. Results

### 2.1. Identification of PLD inducing compounds from drug repurposing collection

To investigate PLD in an experimental setup, we used 5,228 compounds from the repurposing collection inspired by the Broad Repurposing Library (v. 2017)^33^. The compounds were screened at 10 µM in Vero-E6 and A549-ACE2 cells for induction of PLD after 24 h of treatment. At first sight, our observations revealed that certain classes of compounds caused PLD across both cell lines, while others were exclusive to specific cell lines. This suggests a potential cell line dependency in drug-induced PLD (Figure 1). In summary, according to applied criteria described in ( 4.2. Building training and test sets) we observed 155 molecules inducing PLD in A549-ACE2 cells and 197 molecules inducing PLD in Vero-E6 cells. These numbers represent 2.7-3.5 % of the total drug repurposing collection, respectively.

**Figure 1.**
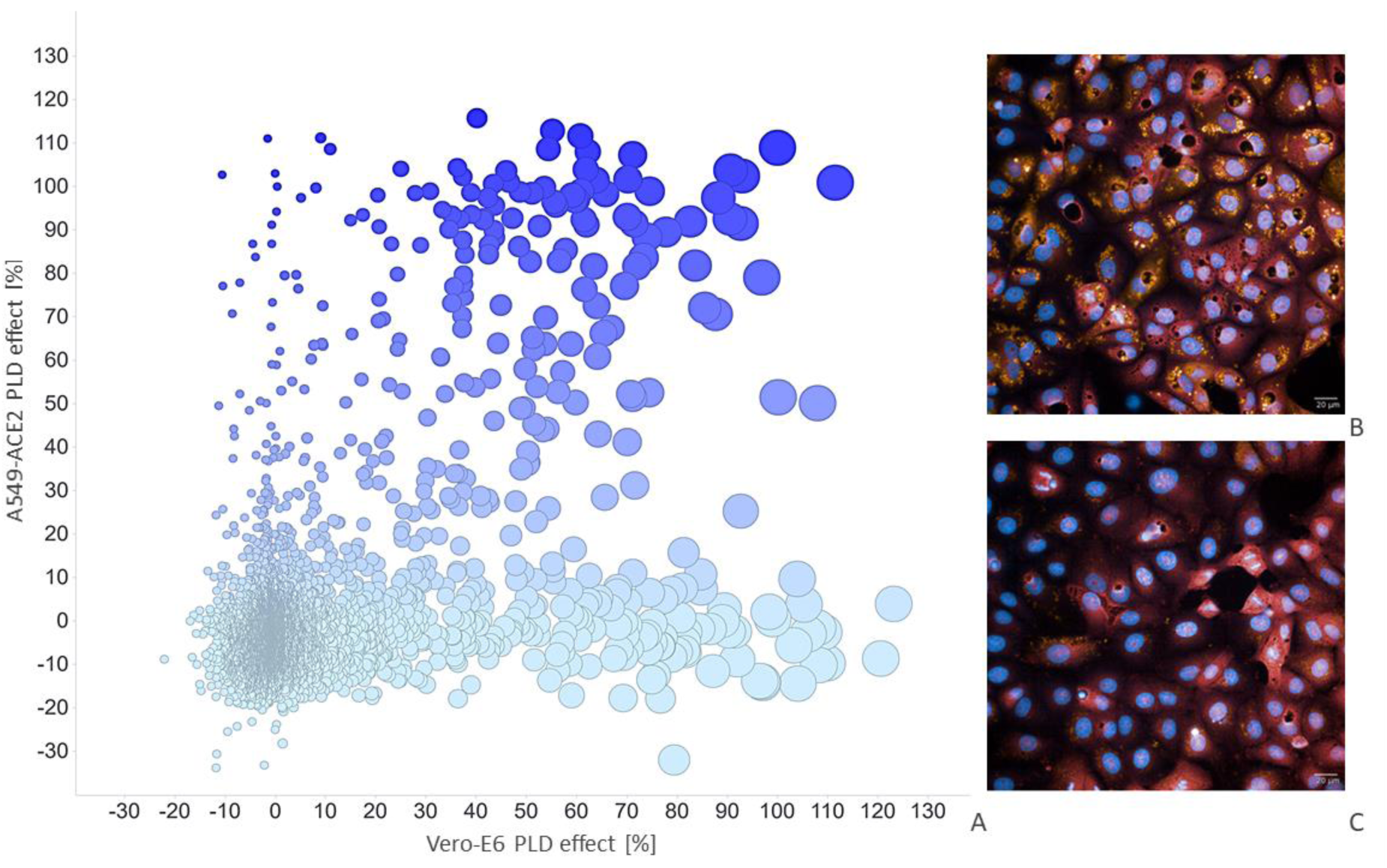
Correlation of PLD in Vero-E6 and A549-ACE2 cells. A: plot showing the correlation of the PLD induction screening results between Vero-E6 cells and A549-ACE2 cells. Data points a marked by color for A549-ACE2 PLD effect and by dot size for Vero-E6 PLD effect. B: PLD induction in Vero-E6 cells using amiodarone 20 µM (positive control for PLD induction). C: Vero-E6 cells incubated with 0.1 v/v % DMSO (negative control for PLD induction). Blue: Nuclei stain, red: cytoplasm stain, orange: PLD detection.

Next, we focused on generating a representative dataset showcasing the PLD effect. For this, we categorized compounds as “active” or “inactive” based on a PLD induction threshold of ≥52% for A549-ACE2 cells and ≥50% for Vero-E6 cells. These different thresholds enabled adjusting for experimental variability across institutes (ITMP and KI) and cell lines. Ultimately, from 5,228 compounds tested, 4,258 compounds with consistent and comparable results across cell lines (i.e. active or inactive in both cell lines) were selected as the representative set and used for ML modeling (Figure 2). After selecting the dataset, we divided the 4,258 compounds into training and testing subsets using an 80:20 split ratio. The chemical structures in the dataset were converted into vector features (through fingerprints) to represent their characteristics, enabling their use as input for the ML model (see Section 4.2, “Building training and test sets” for more details). To ensure the validity of the testing dataset, we carefully analyzed its underlying chemical space prior to model training.

**Figure 2:**
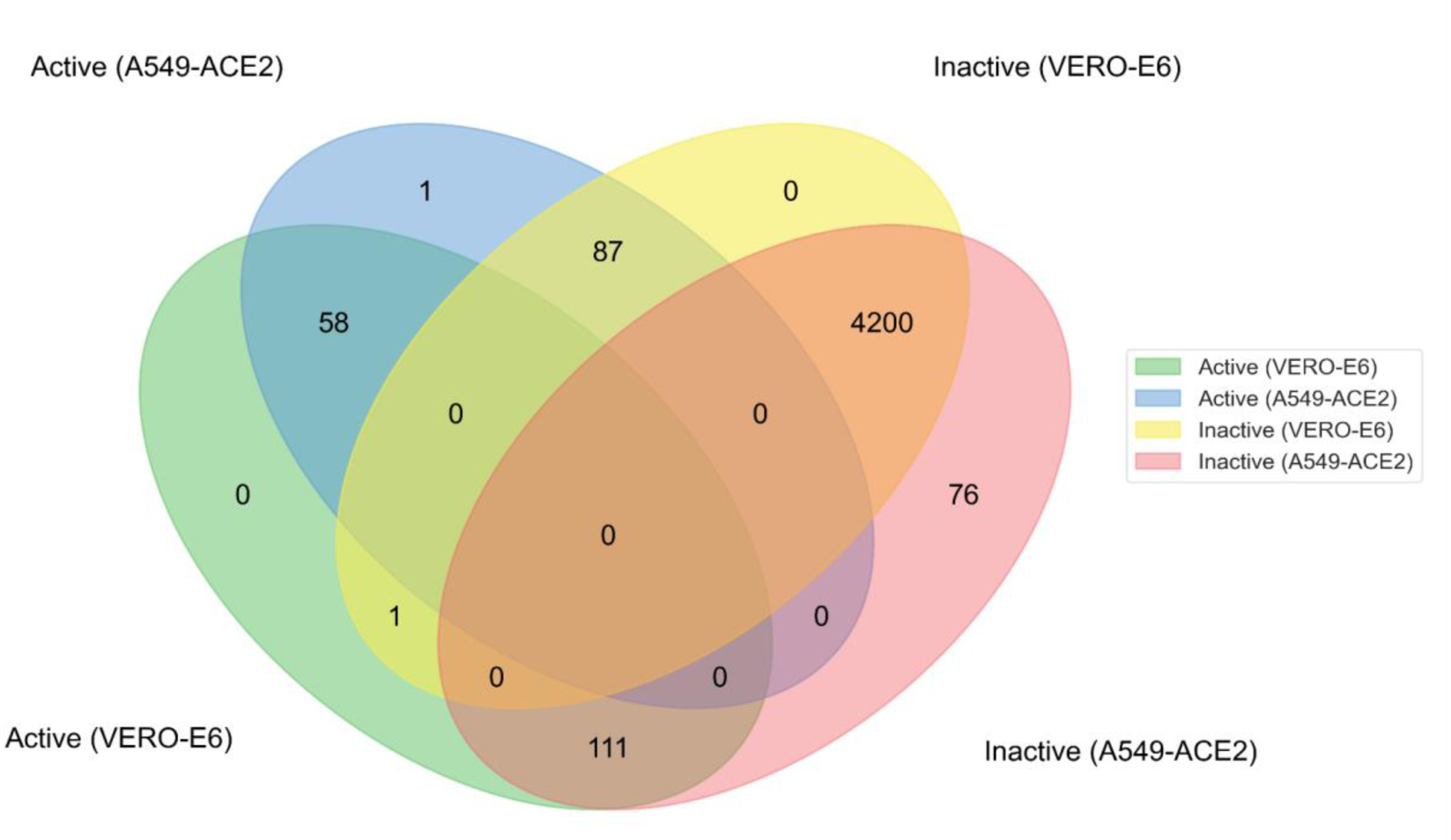
Overlap of the compounds across cell lines. In total, 4,200 compounds found inactive in both screens (Vero-E6 and A549-ACE2) were added to 58 compounds found there active in both screens. The remaining compounds (1,192 compounds) comprise actives or inactives in one of the screens.

To assess the distribution and uniformity of the chemical space across the training and testing sets, we employed a t-SNE (t-distributed Stochastic Neighbor Embedding) plot, as shown in Figure 3. This visualization was critical in confirming that the chemical space modeled through the ErG fingerprinting method, combined with ChemPhys descriptors, was both homogeneous and representative of the broader dataset. Such inspection ensured that the dataset was free from significant biases, which could otherwise compromise the model’s ability to generalize effectively.

**Figure 3:**
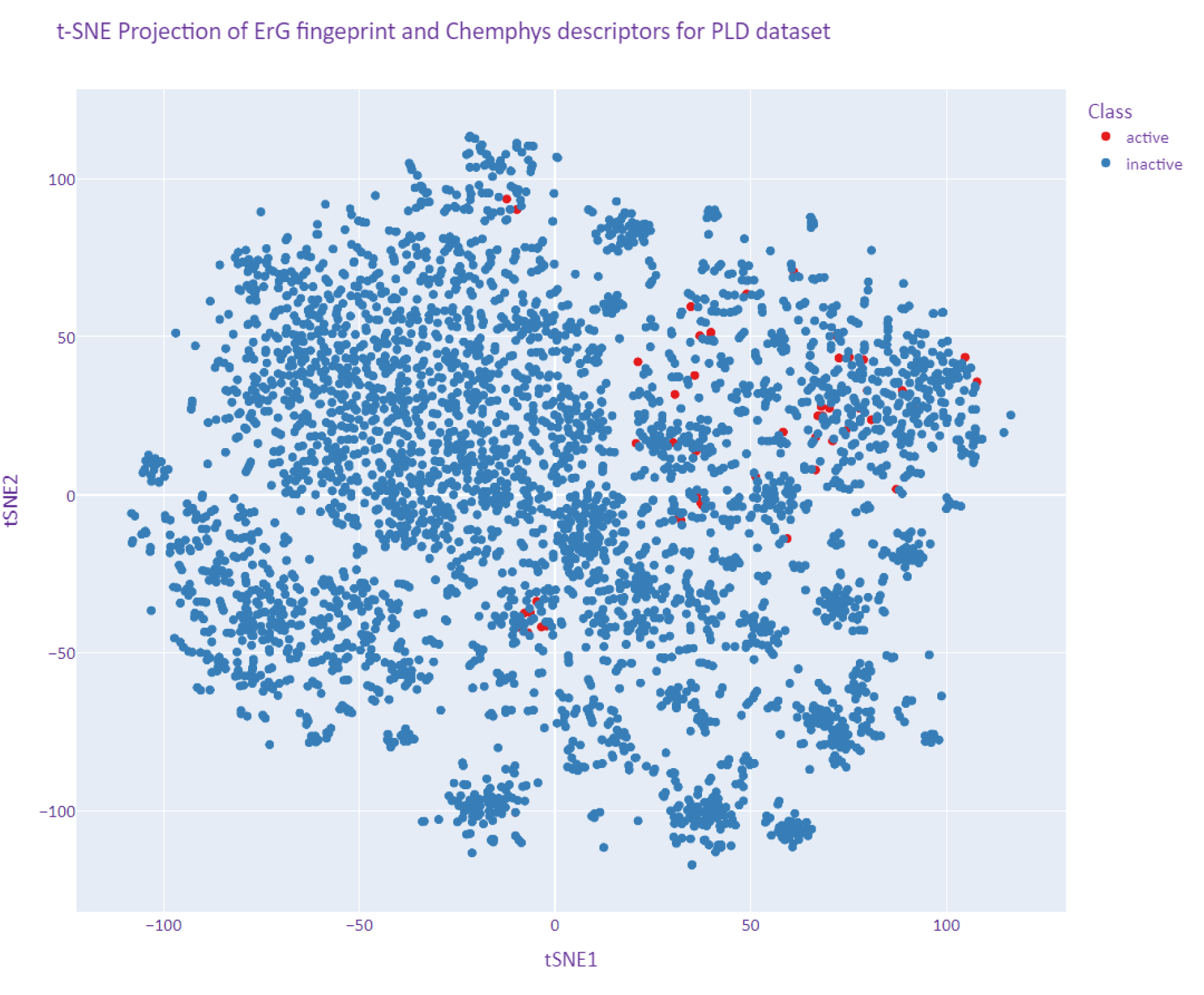
t-SNE plot of the Broad Repurposing library. The dots in blue demonstrate PLD inactive compounds, and those in red highlight PLD active compounds. The homogeneous distribution of actives within the library is evident as it does not show any particular clusterization in the chemical space described by chemical descriptors.

### 2.2. Enhanced performance with Boosted Tree models

Our training approach involved evaluating six machine learning models (Gradient Boosted Tree, Logistic Regression, Neural Network, Decision Tree, Random Forest, and XGBoost) using a 4-fold cross-validation strategy. This method allowed us to systematically assess model performance across different subsets of the data. The results revealed that algorithms based on Boosted Trees performed slightly better than the other models. However, the differences in accuracy were initially overestimated due to a bias in the dataset favoring inactive compounds. When additional performance metrics, such as False Negatives (FN) and False Positives (FP), as reported in Table 1, were considered, the superior performance of XGBoost became more evident. These metrics provided a clearer and more balanced evaluation of the model’s ability to correctly identify active and inactive compounds.

**Table 1:**
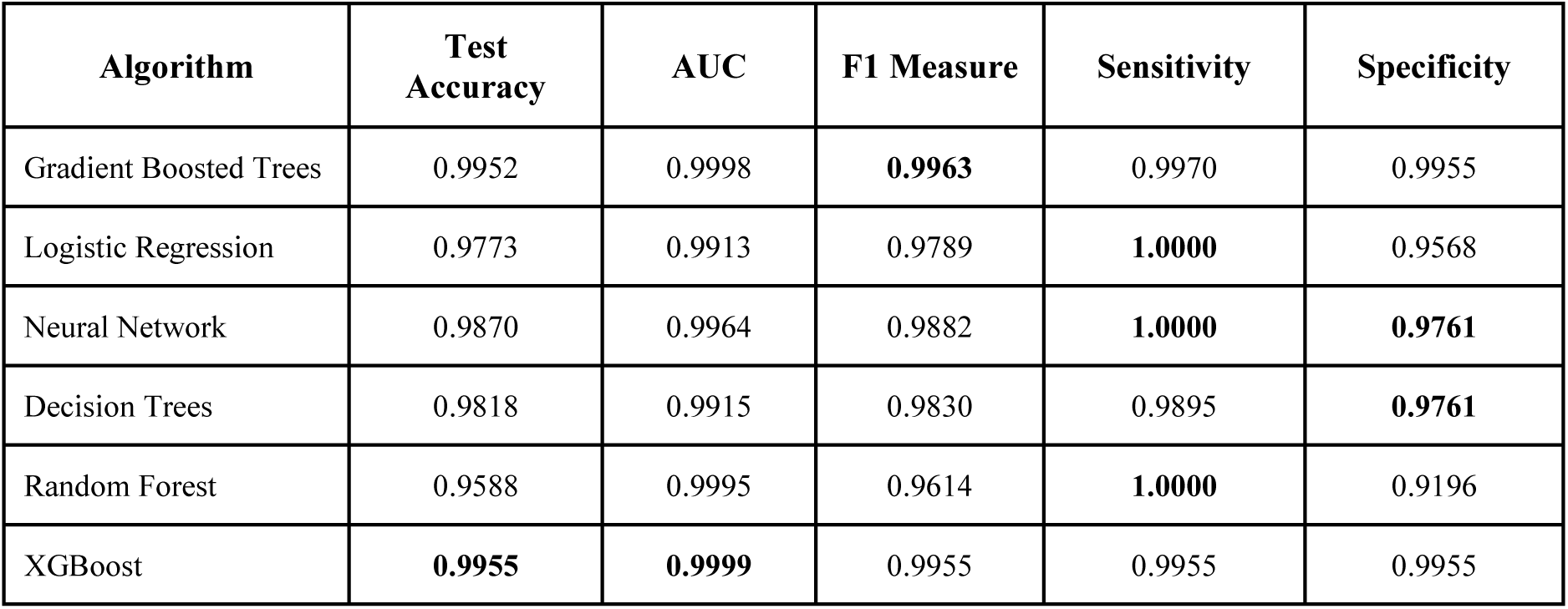
Classification algorithm performances on the test data (N=852). The results shown in the table are the average metric reported for four cross-validation runs. Bolded are the highest values for each metric.

| Algorithm | Test Accuracy | AUC | F1 Measure | Sensitivity | Specificity |
| --- | --- | --- | --- | --- | --- |
| Gradient Boosted Trees | 0.9952 | 0.9998 | **0.9963** | 0.9970 | 0.9955 |
| Logistic Regression | 0.9773 | 0.9913 | 0.9789 | **1.0000** | 0.9568 |
| Neural Network | 0.9870 | 0.9964 | 0.9882 | **1.0000** | **0.9761** |
| Decision Trees | 0.9818 | 0.9915 | 0.9830 | 0.9895 | **0.9761** |
| Random Forest | 0.9588 | 0.9995 | 0.9614 | **1.0000** | 0.9196 |
| XGBoost | **0.9955** | **0.9999** | 0.9955 | 0.9955 | 0.9955 |

**Table 2.**
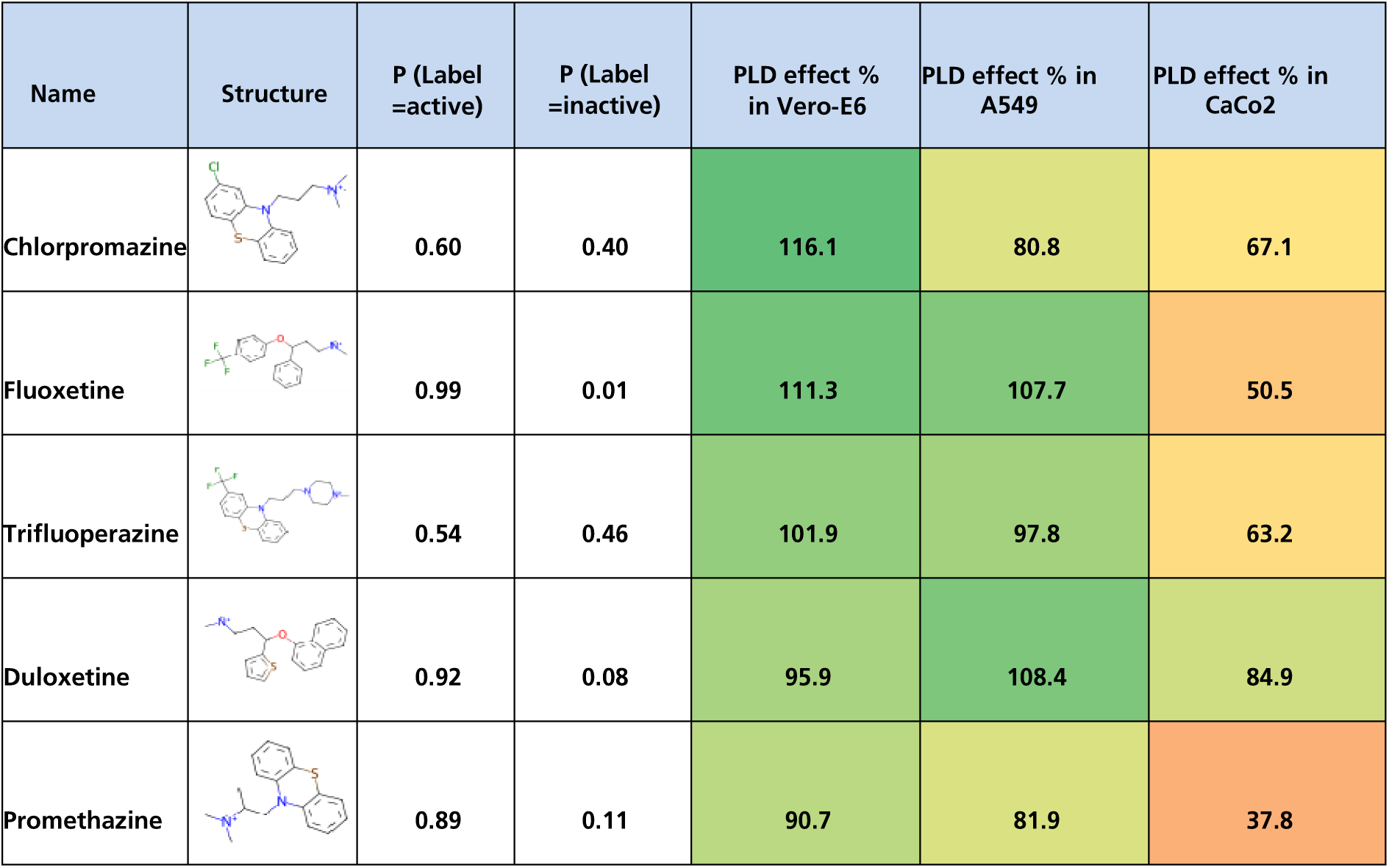
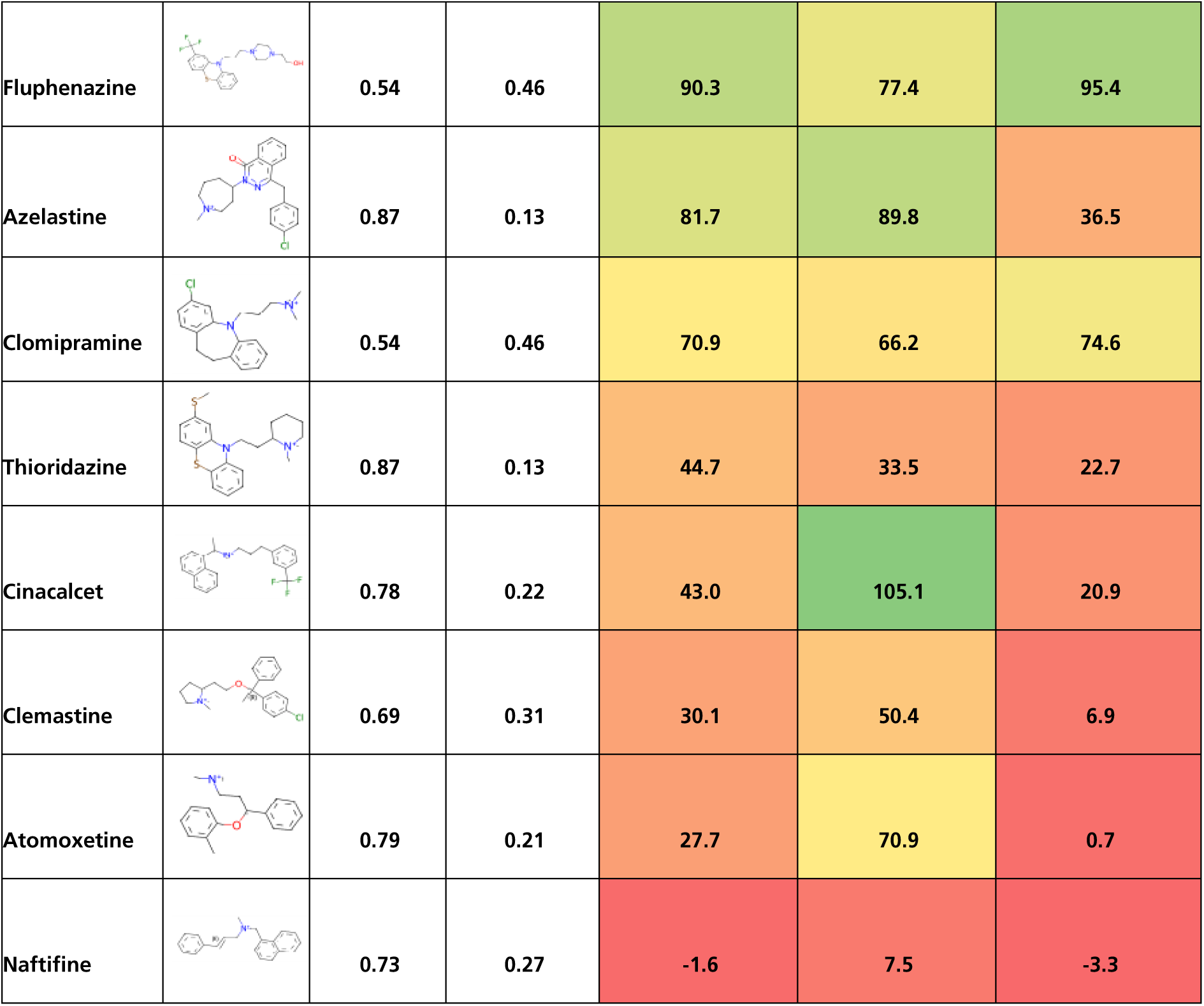
Summary of experimental PLD assay results of ENZO library on three different cell lines measured at 10 µM in-assay concentration.

An analysis of the top 10 most relevant descriptors identified by the model highlights two key observations (Supplementary Figure 1). First, the ChemPhys descriptors appear to be less effective in modeling the problem than initially expected. While characteristics such as cell permeability or solubility, approximated by higher calculated TPSA or logP values, might be anticipated to influence PLD effect, they did not appear to play a significant role in this context. Second, eight out of the ten most relevant features contained stable positively charged atoms at physiological pH, aligning with the known properties of cationic amphiphilic drugs, which are well-documented as PLD inducers ^21^. Additionally, specific molecular structural features, many of which have been validated in previous studies^5^, were instrumental in driving the model’s accurate predictions. For example, several ErG bits describing the distance between aromatic centroids and a positive charge (e.g., Ar_+_d7, Ar_+_d6, Ar_+_d5, Ar_+_d1) ranked first, second, seventh, and tenth, respectively, in the relevance list (Figure 4). Closely related to this finding is the identification of Hf_+_d1 (ranked third), Hf_+_d2 (ranked sixth), and Hf_+_d12 (ranked ninth) among the most relevant descriptors (Supplementary Figure 2). The Hf_+_d1 structural feature corresponds to fragments where isopropyl (iPr), isobutyl (iBu), or lipophilic fatty chains are attached to a nitrogen atom that remains stably charged at physiological pH. This feature frequently includes piperidine-based tertiary or quaternary nitrogen structures. Interestingly, some previously unrecognized structural pharmacophore features also emerged as significant, particularly those representing long-distance interactions. For instance, Hf_Hf_d9 (ranked fifth), +_+_d12 (ranked sixth), and D_Ac_d11 (ranked eighth) have not been reported in prior studies. These novel descriptors suggest new avenues for understanding the molecular basis of the modeled phenomena and highlight the importance of considering unconventional structural features in future analyses. The real value differences for the top 10 features are reported in Supplementary Figure 4 as box plots, together with the relative statistical p-values. Very few non-significant differences were found.

**Figure 4:**
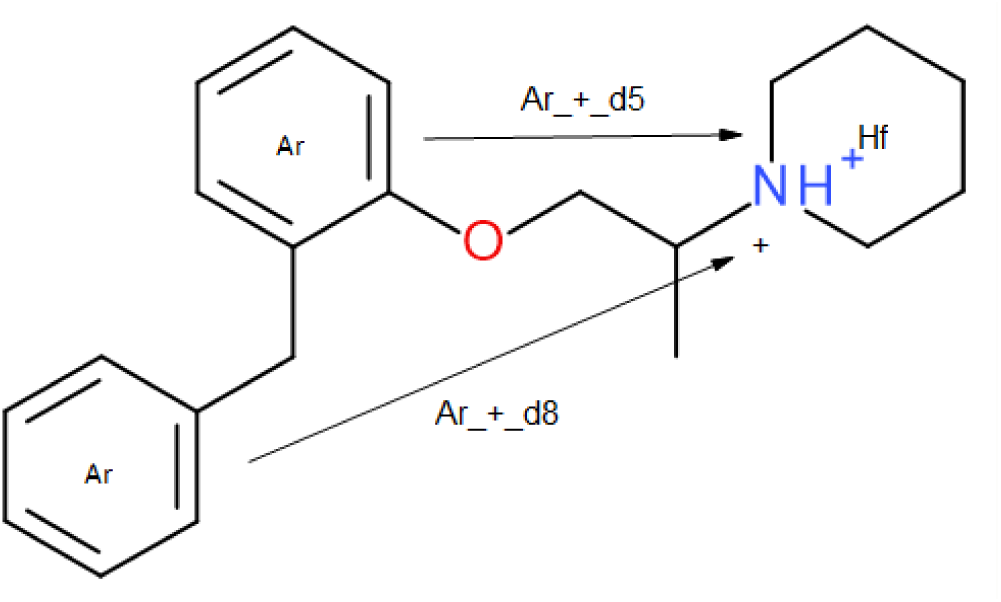
Scheme of Benproperine, an antitussive compound with high PLD propensity, representing how the ErG fingerprint scheme coded the relevant distances.

Finally, we validated the model predicted on an external library. For this purpose, we chose the U.S. Food and Drug Administration (FDA) collection of 774 drugs, referred to here as ENZO^22^. Of the entire collection, 13 compounds (1.7%) were predicted to induce PLD. Compounds labelled as active (P label > 0.5, with P label = probability value to be active with label = 1 against inactives label = 0) were subsequently tested in Vero-E6 cells under the same experimental conditions used in the training set. To assess the model’s transferability to cell lines not included in the training data but frequently used as cellular models especially in phenotypic (antiviral) screens, additional tests were conducted using A549 and CaCo2 cells. At a concentration of 10 µM, 11 out of the 13 predicted compounds (84.6% confirmation rate) demonstrated PLD induction in Vero-E6 cells. Similarly, 12 out of 13 compounds (92.3% confirmation rate) showed PLD activity in A549 cells. In CaCo2 cells, 8 out of the 13 compounds (66.7% confirmation rate) were confirmed to induce PLD.

### 2.3. Correlation of drug-induced PLD with phenotypic screens for SARS-CoV-2 inhibition/Antiviral activities correlations to PLD effects?

The interest in understanding PLD induction and its predictive modeling has recently grown, particularly due to observations that antiviral compound effects against SARS-CoV-2, often associated with PLD induction, have not been consistently replicated *in vivo*^4^ This raises the question of how much PLD might confound screening results, as seen in aggregation issues during SARS-CoV-2 drug screens or other antipathogenic assays^23,24^. In our recent work, we published several phenotypic anti-SARS-CoV-2 screening results, which are now available in public databases (CHEMBL4495565 for Vero-E6 cells and CHEMBL4303101 for CaCo2 cells)^25,26^. Here we use the data to investigate on the drug-induced PLD in our phenotypic anti-SARS-CoV-2 screens. Figure 5 illustrates the correlation between the inhibition of cytopathic effect (CPE) caused by SARS-CoV-2 infection and drug induced PLD observed in these phenotypic screens. For this drug repurposing collection, no significant positive correlation was observed between PLD induction and the CPE inhibition. When comparing assays across cell lines, Vero-E6 cells appeared more restrictive than CaCo2 cells in terms of identifying compounds with antiviral activity. On the other hand, we saw from the test of the ENZO library collection that Vero-E6 cells seam to show higher sensitivity to PLD compared to CaCo2 cells. By correlating PLD induction and CPE inhibition in Vero-E6 cells, 2 out of 35 compounds with inhibition of CPE of >30% also exhibited PLD induction of >50% (5.7 % of the total hits). In CaCo2 cells, 68 out of 956 compounds (7.1% of the total hits) showed inhibition of CPE of >30% and PLD induction of >50%. This observation suggests that although the total number of anti-CPE compounds seems to be very different; the percentage of compounds among anti-CPE that also show PLD is quite similar. These findings highlight a potential confounding factor in anti-SARS-CoV-2 screening programs, as although there is no direct positive correlation, drugs inducing PLD alongside antiviral effects may be falsely identified as hits.

**Figure 5.**
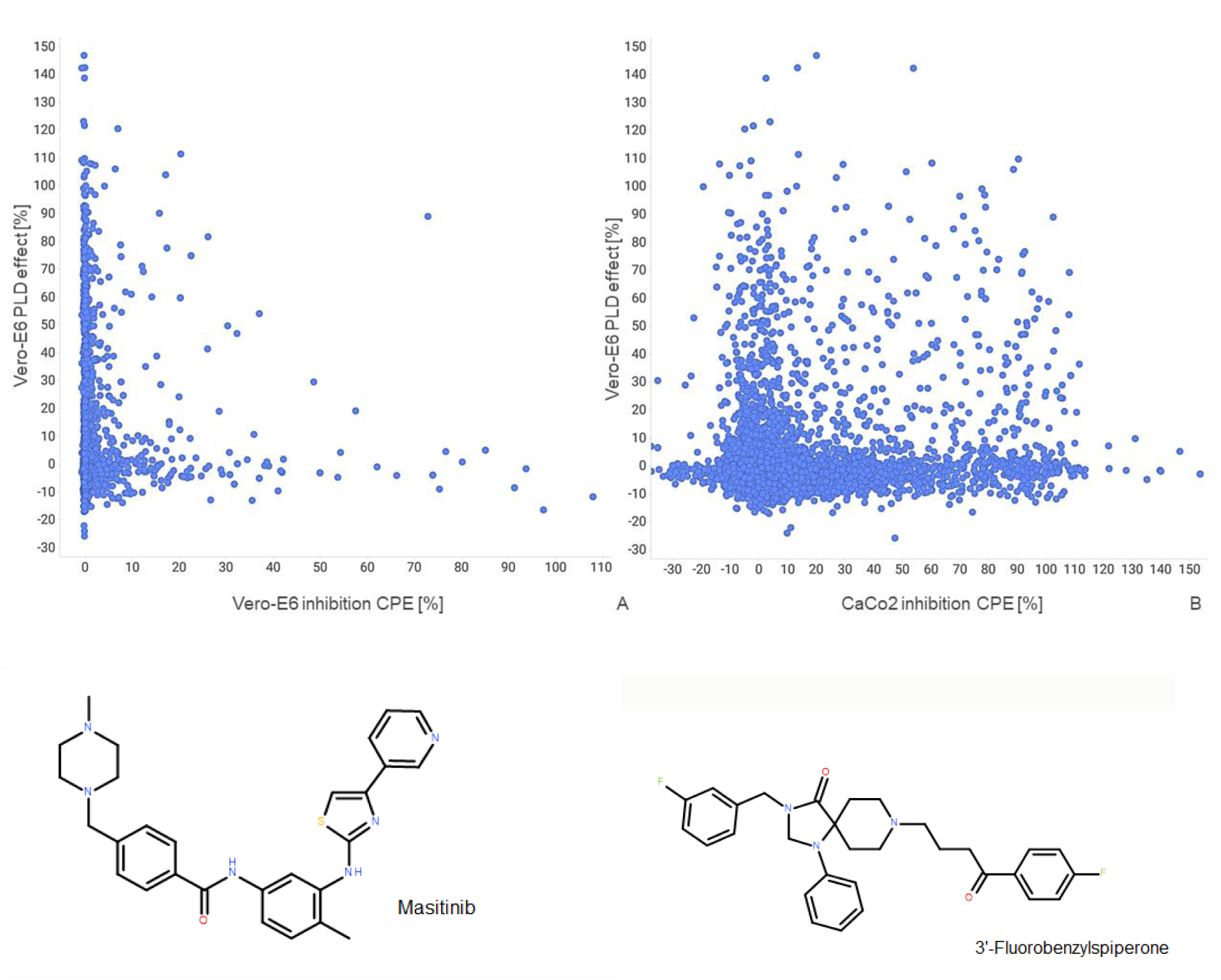
Correlation of PLD induction and SARS-CoV-2 CPE inhibition. A: Scatter plot correlating PLD-induction in Vero-E6 cells with CPE inhibition as reported in CHEMBL4495565 CPE assay on Vero-E6 cells. B: Scatter plot correlating PLD-induction in Vero-E6 cells with CPE inhibition in Caco2 cells as reported in CHEMBL4303101 CPE assay. Bottom: Two examples of active found compounds in PLD induction assay and in Vero-E6 and CaCo2 CPE assay: masitinib and 3’-fluorobenzylspiperone.

The question of how much of the observed antiviral activity can be imputed to PLD induction is certainly legitimate. The presented correlation is based on primary screening results for both PLD and anti-CPE expressed as percentage of positive control measured at 10 µM. A deeper look into drug-induced PLD EC50 and anti-CPE EC50s can reveal a notion, which may improve the discriminant between PLD and anti-CPE in some antiviral screening. From 68 dual active compounds in CaCo2 cells we have selected compounds with strong CPE inhibition (>50 %) and PLD induction (>50 %), including the 2 dual active compounds in Vero-E6. These compounds were tested in dose-response in Vero-E6 cells to determine PLD EC50 and correlated with available anti-SARS-CoV2 CPE EC50 data (Supplementary Table 1). From the dataset we identified 9 compounds for which both anti-CPE and PLD EC50 data were available: NNC-55-0396, lidoflazine, PD-161570, U-18666A, mefloquine hydrochloride, casin, amodiaquine dihydrochloride, 3’-fluorobenzyl spiperone and masitinib (AB1010). We observed that for these compounds the anti-CPE EC50 was higher than then the calculated PLD EC50, meaning that the phenotypic antiviral effect was observed upon the onset of PLD induction. For 16 compounds we also observed that the highest concentration of 20 µM results in onset of a toxic effect, shown as negative PLD % effect values. Although the dataset represents a relatively low sampling size, the calculated EC50 revealed no drug-induced PLD activity in low nM level. Most compounds showed an EC50 between 1-3 µM, suggesting a potential activity range of this drug-repurposing collection.

## 3. Discussion

Phospholipidosis is a long-known storage disorder of accumulated phospholipids in lysosomes, with first research results published in early 1948s. Among classical counter screenings used, PLD might play a secondary role when compared to PgP, hERG or specific selectivity screening. However, in certain fields, like antimicrobial and antiviral efforts, PLD plays an important role either in early infection (cell penetration) or late pathogen maturation mechanisms including cell exit pathways. In some cases, we witnessed compounds with proven antimicrobial and antiviral activity, which in reality turned out to be interfering with host mechanisms of lipid assembly or trafficking. In recent years, the pharma industry has raised interest in testing and predicting this phenomenon, not least because antiviral effects associated with drug-induced PLD in SARS-CoV2 research have not been consistently replicated *in vivo,* but also due to increasing focus of the Food and Drug Administration (US FDA) on potential toxicity of drug-induced PLD. Drug-induced PLD is part of the FDA Adverse Event Reporting System (FAERS), with 60 reported severe cases of drug side effects in the last 5 years, among them one death case^27^. Therefore, we analyzed the largest reported data set of repurposed drugs, that includes 5,228 clinical compounds and drug candidates for PLD induction.

With a rigorous approach in merging different screening results and removing from the training dataset the molecules, actives as well as inactives, inconsistently labelled in different cell lines, we developed a machine-learning classification method, that allowed us to successfully predict external repurposing collections like ENZO, opening up a more general perspective for filtering library collections with dubious or certain PLD-inducing pharmacophoric chemical features. Among the chemical features collected by the model, the already known amphiphilic properties involving stable charged nitrogens in diverse distances from hydrophobic groups, are certainly the most evident. However, within the top 10 most influential molecular features there are important features described in the literature: for instance, the appearance of Hf_d1 and Hf_d2 showed evidently that the stable positive charge needs to be very close (one bond or two maximum) to lipophilic atoms. Such typical arrangements are di-isopropyl or di-butyl amino groups, which are known to be stably positively charged but easily water-de-solvatable. Totally novel and unprecedented molecular feature is D_D_12, i.e. a long distance (12 bonds) pair of hydrogen-bond donors, which, however, looks not so prominent in favor of PLD active molecules as reported in boxplot plots.

From experimental evaluation we observed that depending on the readout 2.7-3.5 % of the total 5,228 compounds in the drug repurposing collection are prone to induce PLD. Also, the prediction for PLD inducing compounds in the ENZO collection resulted in 1.7 % predicted as active. In contrast, our analysis of different library collections like LifeChem^28^, 50k diversity set using our prediction model resulted in 0.09 % PLD-active hits. The results suggest a possible higher accumulation of different PLD-active compounds among the repurposed drugs and/or a more effective and precise filtering of the chemical space in commercial library design.

The application of Vero-E6 and A549-ACE2 in PLD screening already showed a difference in sensitivity of cell lines for PLD. The observation was confirmed during the ENZO compound evaluation in Vero-E6, CaCo2 and A459-ACE2 cells. In addition, with EC50 determination we observed PLD can as well be a toxic phenomenon when compound doses are increased > 20 µM. Whereby, the effect of PLD itself, in most cases, was observed between 1-10 µM compound concentration in assay. In our analysis of the impact of drug-induced PLD on anti-SARS-CoV-2 compound effect, we saw no evident positive correlation between PLD induction and CPE inhibition in Vero-E6 and CaCo2 cells using the repurposing collection. It is not yet known whether the reported infection susceptibility is mechanistically linked to PLD susceptibility. However, for Vero-E6 and CaCo2 cells the two susceptibilities appear to be correlated. For instance, when measuring SARS-CoV-2 infection susceptibility Caco-2 cells were not highly susceptible to SARS-CoV-2 infection (25% infection rate), while Vero-E6 cells showed 77% infection rate^29^. Also, during the ENZO compound evaluation Vero-E6 cells showed higher PLD sensitivity (later confirmed for the whole ENZO library collection, data not shown).

What was not explored in this study, but certainly will have an impact on PLD induction, is the duration of exposure to the compound. The combination of cell line sensitivity, screening concentration and exposure time will certainly impact how much PLD is confounding the desired compound effect or reveal PLD as a side effect in phenotypic screening for a compound.

The question lingering in the background, whether induction of PLD can be leveraged in antiviral/antibacterial treatments or should be excluded from screening hit identification efforts remains unanswered yet. Our study provides to the scientific community a comprehensive dataset for analysis of PLD induction by repurposed drugs, that together with the ML model allows a deeper exploration of chemical features of PLD induction and of impact on in-cell compound activity in future.

## 4. Methods

### 4.1. Experimental protocol for PLD

PLD induction was assessed at both the Fraunhofer Institute for Translational Medicine and Pharmacology (ITMP) and the Karolinska Institute (KI) using their respective drug repurposing collections^30,31^. These collections were mainly developed with design features inspired by the Broad Repurposing collection and acquired in parallel from the same library provider^32,33^. We outline two protocols below, with ITMP conducting studies on Vero-E6 cells and KI on A549-ACE2 cells.

At ITMP, our experimental approach involves using Thermo’s high-throughput screening (HTS)-optimized kit, which offers uniformity, reproducibility, and a narrow emission range suitable for multiplexing. To measure drug-induced PLD in Vero-E6 cells, we used the HCS LipidTOX™ Red Phospholipidosis Detection Reagent (Invitrogen™, #H34351), which is non-toxic to cell growth and remains well-retained after aldehyde fixation. In this setup, compounds (at 10 µM concentration) and controls (amiodarone 10 µM concentration; DMSO 0.1 v/v %) were added to clear bottom, 384-well plates (PhenoPlate 384-well, black, optically clear flat-bottom, tissue-culture treated, Revvity #6057302) using the Echo Liquid Handler (Labcyte). Vero-E6 cells were cultured in a complete medium: Dulbecco’s Modified Eagle Medium (DMEM) supplemented with 10% FBS, 6 mM L-glutamine, and 1% penicillin/streptomycin. Cells were harvested at 80% confluence from a T175 cm² flask by washing once with 10 mL of RT PBS, followed by incubation with 5 mL of Trypsin 0.05% / EDTA 0.02% for 5 minutes. Cells were then resuspended in 10 mL of prewarmed culture medium. For plating, Vero-E6 cells were diluted to a concentration of 500,000 cells/mL in cell culture medium supplemented with 1:1000 dilution of HCS LipidTOX™ Red Phospholipidosis Detection Reagent (1000X). A volume of 20 µL of this cell suspension was added to each well of the 384-well plate, and cells were incubated for 24 hours at 37 °C in a 5% CO₂ environment. For fixation, 20 µL of 8% formaldehyde solution was added to each well and incubated for 30 minutes at room temperature. After incubation, the fixation buffer was then removed, and cells were washed with 50 µL/well of PBS. Cells were stained for 45 minutes with 20 µL/well of a solution containing CellMask™ Deep Red stain (1:3,000, Invitrogen #H32721) and Hoechst 33258 (1:10,000, Merck/Sigma-Aldrich #94403) diluted in PBS. Following staining, the solution was removed, and cells were washed three times with 50 µL/well of PBS. Image acquisition was performed using the automated Operetta Phenix High Content Screening system (Revvity). Image analysis was performed using Columbus 2.9 image analysis software (PerkinElmer Inc) (Supplementary Figure 3). Results were normalized against the positive control (amiodarone at 10 µM, set at 100% PLD induction) and the negative control (DMSO at 0.1% v/v, set at 0% PLD induction). Compounds with >50% PLD induction were classified as primary hits.

In a parallel effort at KI, assay-ready plates were prepared using the Echo (Labcyte) and stored in the freezer (−20 °C) until use. Plates with pre-spotted compounds and controls were retrieved and thawed at room temperature for 30 minutes before seal removal. A549-ACE2 cells were cultured in a complete medium: DMEM/F-12 with GlutaMAX™ Supplement (Fisher Scientific, 31331028), 10% FBS heat inactivated (Gibco, Fisher Scientific, 10500064) and 1% penicillin/streptomycin (Cytiva, Nordic Biolabs, SV30010) and 1x NEAA (Cytiva, Nordic Biolabs, SH30238.01). Cells cultured in T175 flasks and reaching 70–90% confluency were detached with trypsin and diluted to 1.3 x 10⁵cells/mL in media. The LipidTOX™ Green reagent (ThermoFisher, # H34475) was diluted in the cell suspension to a final dilution of 1:1000. This cell mixture was dispensed at 30 µL (4000 cells/well) into the pre-spotted plates (PhenoPlate 384-well, black, optically clear flat-bottom, tissue-culture treated, Revvity #6057302) using a Multidrop dispenser. Plates were stacked, covered with wet paper towels, and incubated in a plastic box inside a cell incubator. After 24 hours, cells were fixed by adding 10 µL of 16% PFA (ThermoFisher, #28908) to each well and incubation for 20 minutes at room temperature. The cells were washed three times with 80 µL of PBS using an Aquamax 4000 plate washer (Molecular Devices) and then incubated with 30 µL of a dye mix containing Hoechst 33342 (2 µg/mL, Thermo Fisher, #C10046) and CellMask™ Deep Red (1:20,000, Thermo Fisher, #C10046) in PBS for 1 hour at 37 °C. The plates were washed three additional times with 80µl PBS and stored in 40 µL PBS at 4 °C until imaging, after pre-warming for 10 minutes at room temperature. Images were captured using ImageXpress Micro XLS Widefield High-Content Analysis System. Four fields of view were captured per well using a ×10 Plan Fluor 0.3 NA objective. Images were saved as 16-bit grayscale TIFF files, along with metadata. Image analysis was performed using CellProfiler^34^. For each well, cells were counted from nuclei staining, and cytoplasm was defined from CellMask staining. With in cytoplasm, the number of phospholipid spots were counted from the LipidTox Green staining. The average number of spots per cell per well was used to indicate induced phospholipidosis. Results were normalized against the positive control (tamoxifen at 10 µM, set at 100% PLD induction) and the negative control (DMSO at 0.1% v/v, set at 0% PLD induction).

### 4.2. Building training and test sets

The Broad Repurposing collection v2017 present in both sites, comprising 5,632 compounds tested on both A549-ACE2 and Vero-E6 cells, formed the basis for our training set (https://repo-hub.broadinstitute.org/repurposing#download-data). We converted the compounds and their corresponding PLD induction results into a machine-readable, interpretable format. Due to variations in the PLD induction readouts between the two cell lines (see Section 4.1 “Experimental protocol for PLD”), we first filtered for compounds showing consistent results across both cell lines. Compounds were categorized as “active” or “inactive” based on a PLD induction threshold of ≥52% for A549-ACE2 cells and ≥50% for Vero-E6 cells, adjusting for experimental variability across institutes and cell lines. Ultimately, 4,358 compounds with consistent results across both cell lines were selected for ML modeling. Their structural information was first processed by desalting and calculating their dominant charge states, then converted from SMILES structural representations to 2D fingerprint vectors to serve as ML model inputs. Using the RDKit library^35^, we encoded the compounds into Extended-Reduced Graph (ErG) ^36^ and 28 “classical” physico-chemical (ChemPhys) descriptors. The ErG fingerprint consists of 315 bits summarizing the pharmacophoric profile of each compound, while the ChemPhys fingerprint consists of 28 bits that represent the basic molecular properties of a compound, including topological polar surface area (TPSA), molecular weight (MW), logP, and other structural count-based descriptors. The selection of these fingerprints was driven by their interpretability and transparency, enhancing our understanding of the mechanics behind the ML model prediction. For model training, these fingerprints, combined with the PLD induction labels (active/inactive), were used as an input matrix to predict compound activity. We split the dataset into training and testing sets, using an 80-20 ratio. We then examined the distribution of PLD induction labels (active/inactive) within the training subset and observed a strong imbalance, with active compounds representing less than 2% and inactive compounds over 98%. To address this imbalance, we applied the Synthetic Minority Over-sampling Technique (SMOTE) method to generate synthetic samples, achieving an even distribution between the two labels^37^.

### 4.3. Training machine learning models

To identify the optimal machine learning (ML) model for our application, we began by training six classical ML algorithms using the AutoML node in KNIME^38^ (Figure 6). These algorithms included Logistic Regression (LR), a perceptron-based Neural Network (NN), Gradient Boosted Trees (GB), Decision Tree (DT), Random Forest (RF), and eXtreme Gradient Boosting (XGBoost). The SMOTE-balanced training subset served as the input for this group of models. To ensure the robustness of the trained model, we performed 4-fold cross-validation, leaving out 10% of the data during each trial. The AutoML approach enabled us to identify the best-performing model from this cohort. In addition to the KNIME computational approach, we implemented a Python-based version to train and optimize the KNIME best-performing model (i.e., XGboost classifier). For XGBoost hyperparameter optimization, we leveraged on a refined GridSearch strategy provided by the scikit-learn library. The parameters used in the grid hyperparameter strategy include: “n_estimators” at 700, “learning rate” at 0.025 and 0.05, “max depth” at 8 and 10, “gamma” at 0 and 1, “colsample bytree” at 0.5 and “min_child_weight” with values 2, 3, and 5.

**Figure 6:**
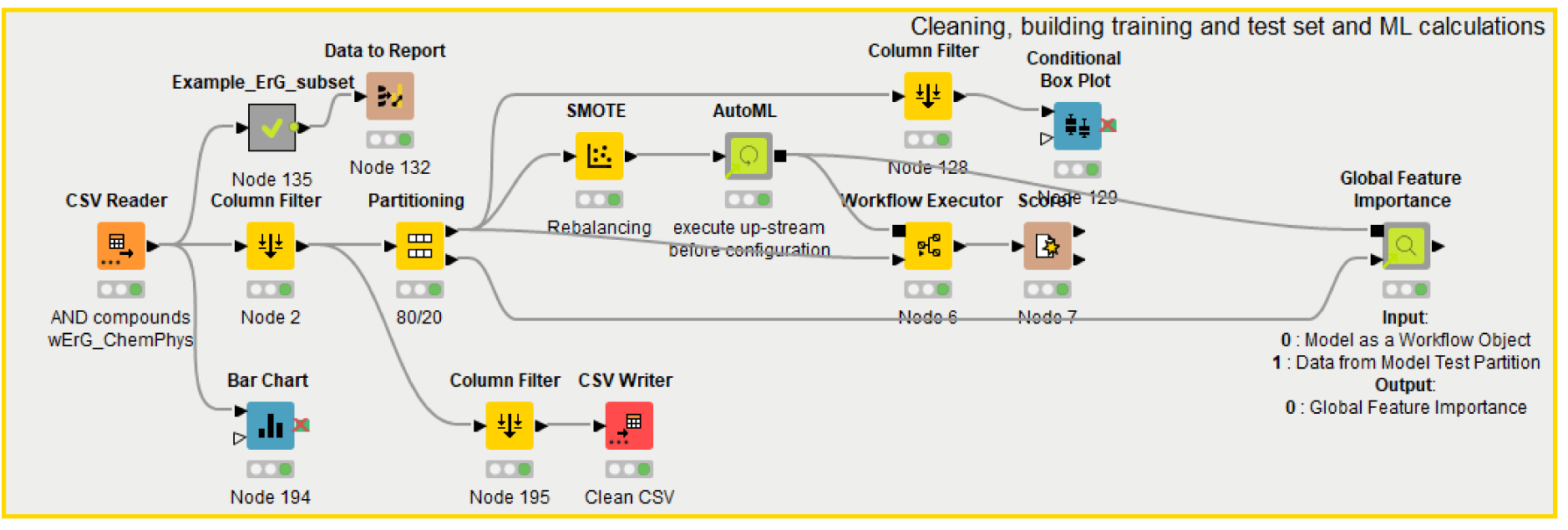
KNIME Workflow used for ML development. This workflow begins with a CSV Reader node to load the training and testing datasets, which include ErG fingerprints and physicochemical descriptors. The central AutoML node then trains a selection of models, and for the top-performing models, feature importance is extracted using the Global Feature Importance node.

In addition to model training and optimization, we conducted a feature importance analysis to identify the key chemo-physical properties that the models used to distinguish between active and inactive compounds in PLD induction. This was achieved using the Global Feature Importance node in KNIME and the feature importance function within the XGBoost Python package ^39^. To assess model performance, we employed confusion matrices and Cohen’s kappa score, allowing us to evaluate and compare model accuracy and reliability. Alongside these metrics, we also report the F1 measure, sensitivity, and specificity as calculated below:

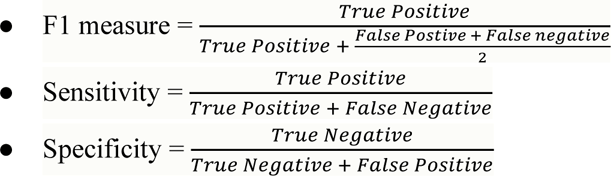

### 4.4. Implementation and data availability

The source code and data files used in this manuscript can be found on resource-specific repositories: code on Github (https://github.com/Fraunhofer-ITMP/PLD) and KNIMEHub (https://hub.knime.com/fraunhoferitmp/spaces/Public/Remedi4All/Phospholipidosis_v5~m6rnKt_4iYtDI1yt/current-state), cellular imaging data will be uploaded on Image Data Repository (IDR) and the models deployed on a public interface on the SERVE (https://serve.scilifelab.se/collections/remedi4all/) instance for community use. We will upload the PLD experimental results data in ChEMBL v36 (expected to be released next year 2026), in the meantime, screening data used in this manuscript is attached as a supplementary file. Both experimental protocols are outlined in detail in a standard operation procedure (SOP) recipe on Zenodo at https://doi.org/10.5281/zenodo.13880869.

## Supporting information

Supplemental Table Screening Data

## Acknowledgments

We would like to thank the authors of the resources used in our work for making their datasets available to the scientific community. We thank Oscar Fernandez-Capetillo Laboratory, Karolinska Institutet, SciLifeLab for generously providing A549-ACE2 cells. We would like to acknowledge Leonie Von Berlin for their feedback in improving the manuscript. We would also like to thank the anonymous reviewers for their comments and suggestions for improving the readability of the manuscript.

## Author contribution

Maria Kuzikov: Conceptualization, Investigation, Data curation, Writing – original draft, Visualization and Validation; Yojana Gadiya: Conceptualization, Methodology, Writing - Original draft preparation and Visualization; Andrea Zaliani: Conceptualization, Supervision, Methodology, Writing - Original draft preparation, Data curation and Visualization; Jeanette Reinshagen: Conceptualization, Methodology, Data curation Writing - Original draft preparation Johanna Huchting: Conceptualization, Methodology, Writing - Original draft preparation Conceptualization, Writing - Reviewing and Editing; Philip Gribbon: Writing - Reviewing and Editing; Adelinn Kalman: Methodology, Investigation and Formal analysis; Kun Qian: Methodology, Investigation and Formal analysis; Hanna Axelsson: Methodology, Investigation and Formal analysis; Marianna Tampere: Methodology, Investigation and Formal analysis; Päivi Östling: Methodology, Investigation and Formal analysis; Brinton Seashore-Ludlow: Methodology, Investigation, Formal analysis and Writing - Reviewing and Editing

## Supplementary material

**Supplementary Figure 1:**
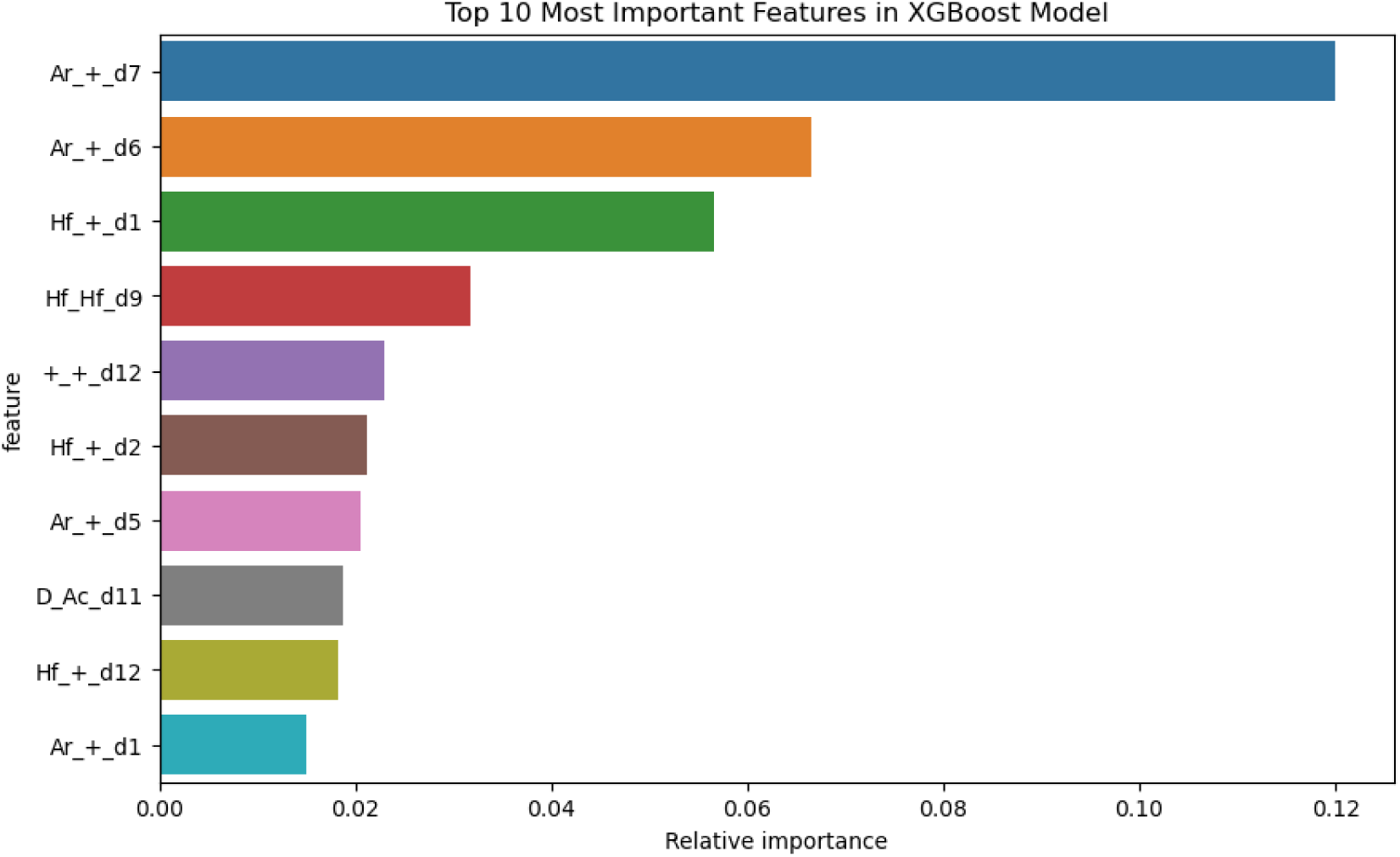
Bar chart reporting the top 10 most influential descriptors for the XGBoost model. The top features shown in this figure are the ErG annotations. In this notation, for instance, a descriptor like “Ar_+_d7” means that bits are set to 1 when an aromatic centroid lies seven bonds distant from a positive charge. Hf identifies a hydrophobic group (a collection of at least three carbon atoms), while “D” and “Ac” identify hydrogen-bond donors and acceptors, respectively.

**Supplementary Figure 2:**
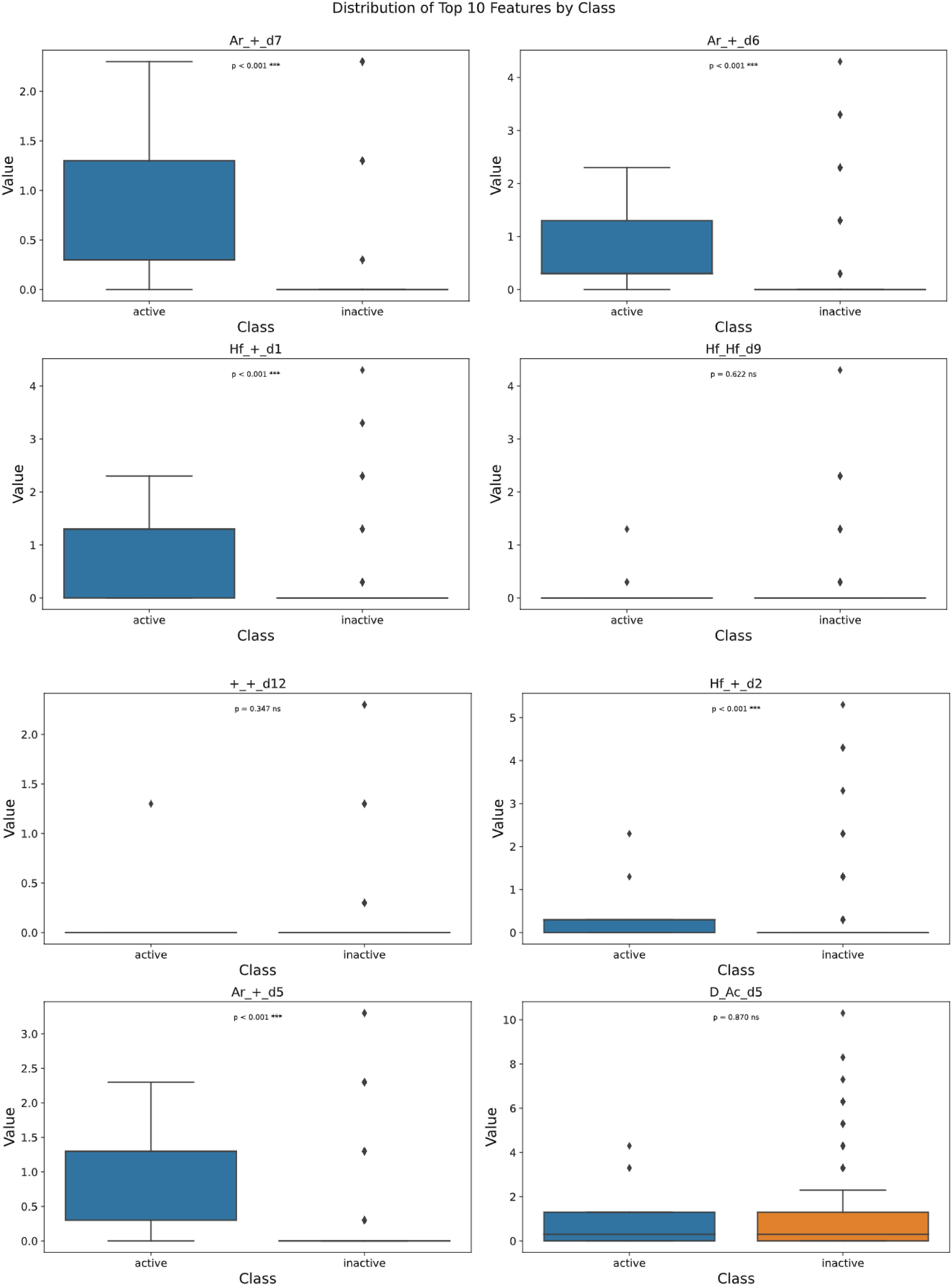
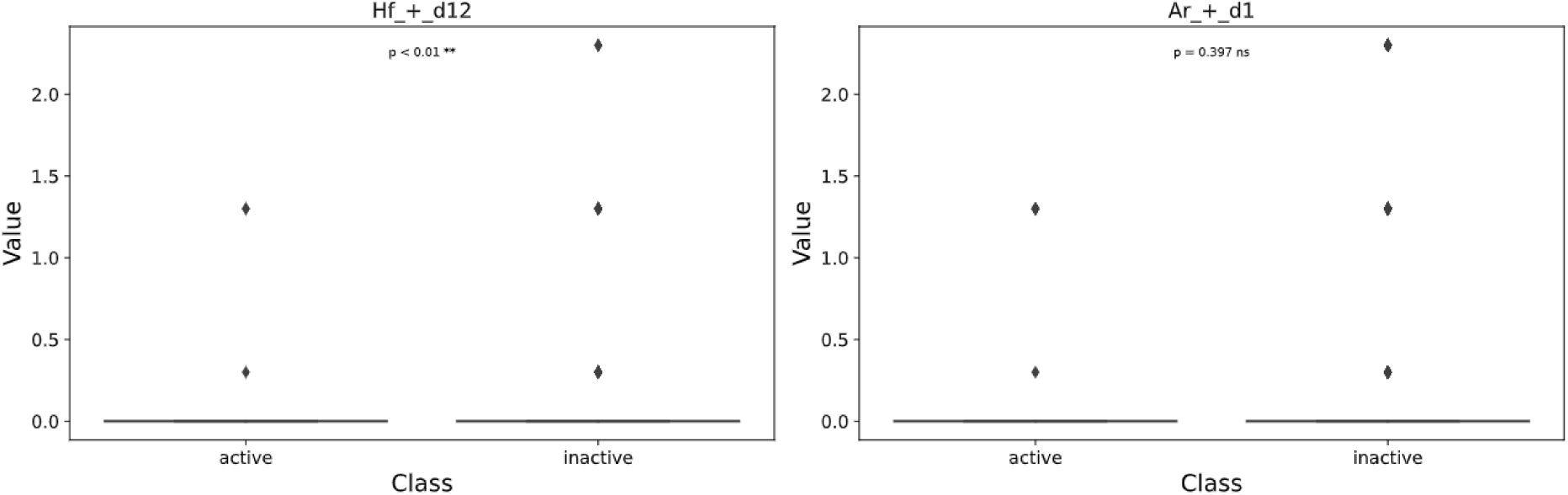
Distribution of the top10 most influential ErG features across the two classes with enhanced box plots. To understand the distribution across the two classes namely Active and Inactive in the original dataset. In almost all features chosen by the XGBoost model, Actives show higher values than Inactives. Significance p-values are given.

**Supplementary Figure 3:**
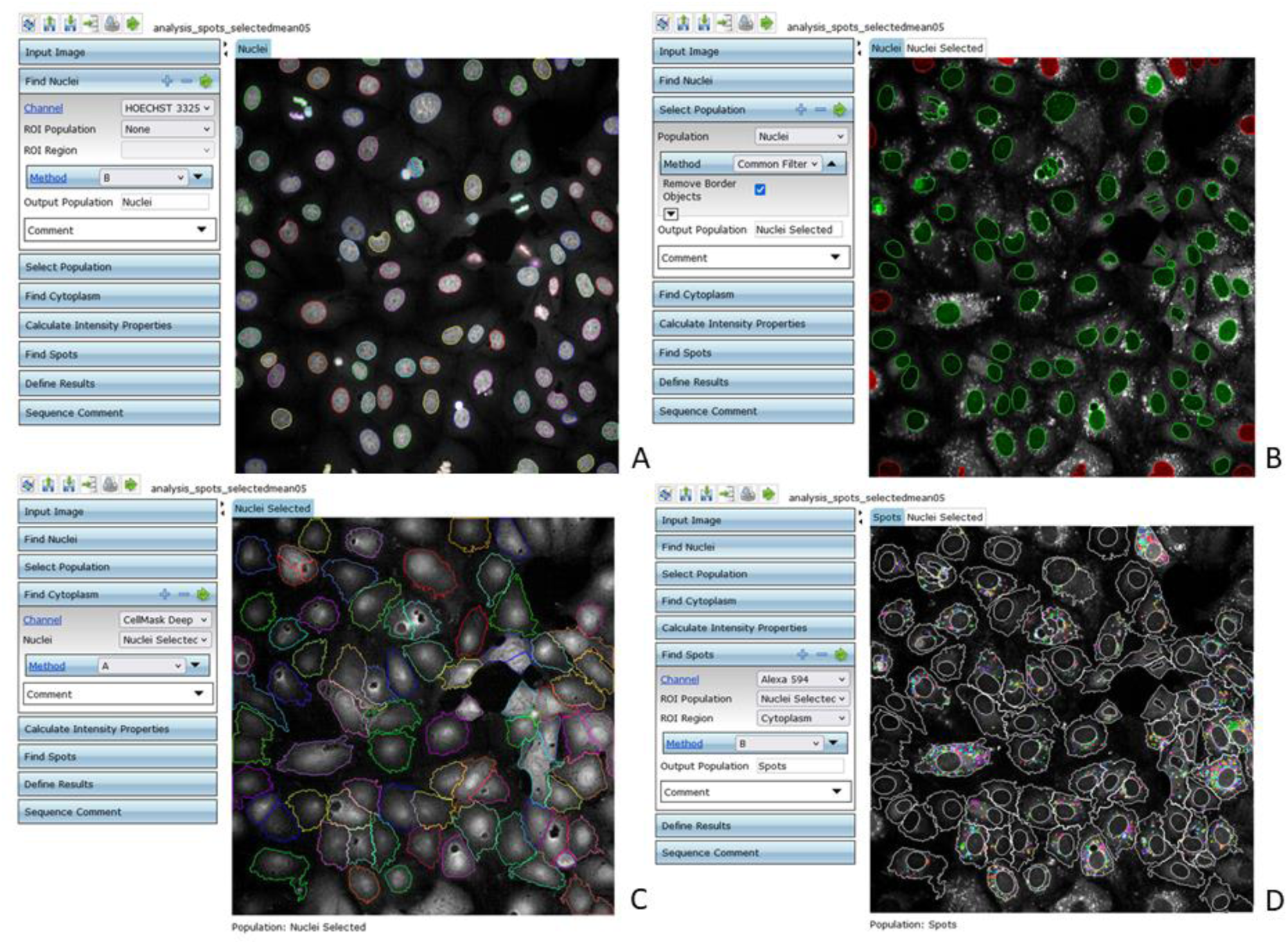
Workflow of DIPL data analysis in Columbus Image Analysis Software. Nuclei identification (A) Removement of border objects (nuclei are completely displayed) = “selected population” (B) Cytoplasm identification in the selected population (C) Calculation of DIPL signal in the cytoplasm: Identification of spots (D).

**Supplementary Table 1.**
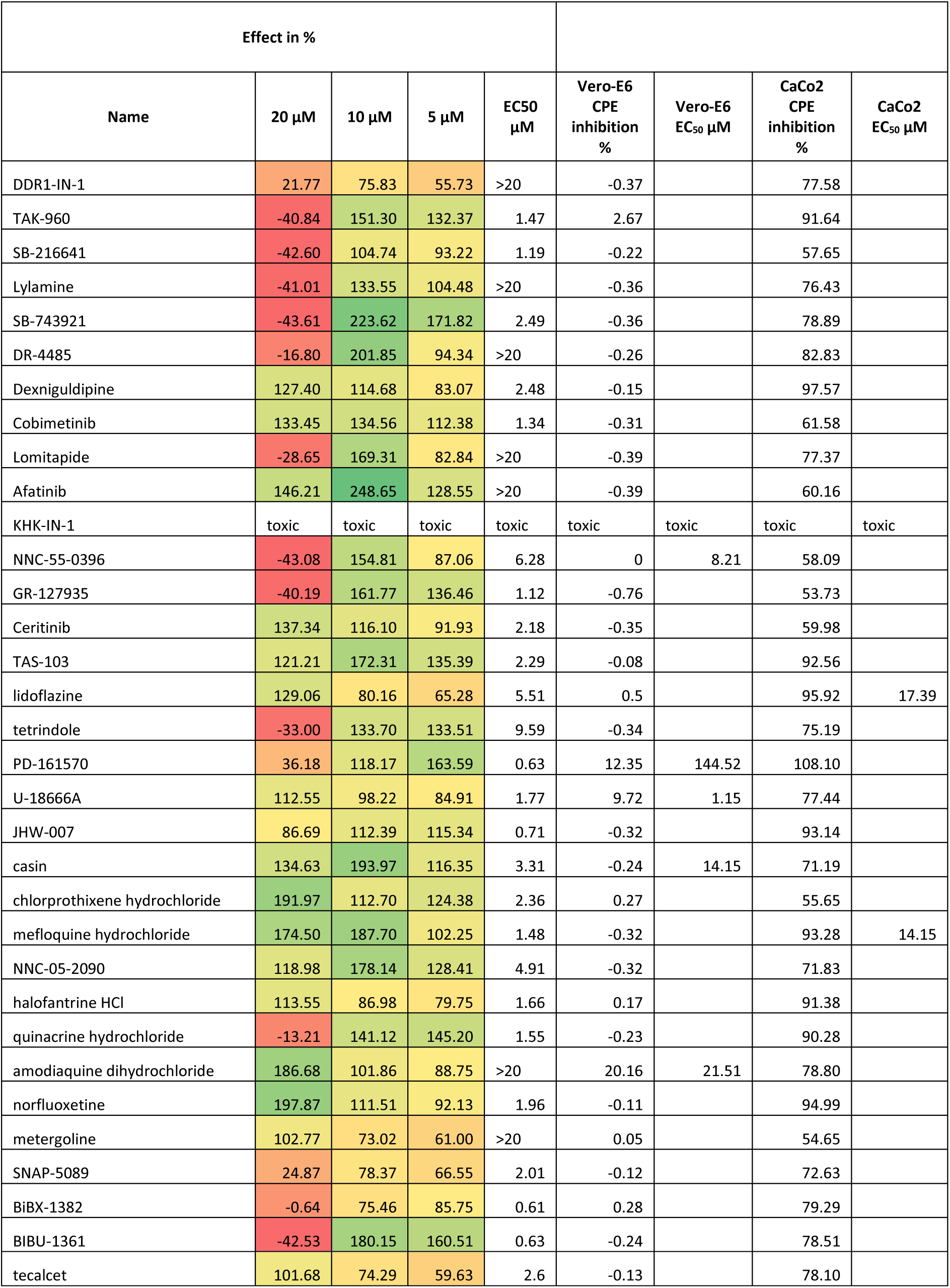
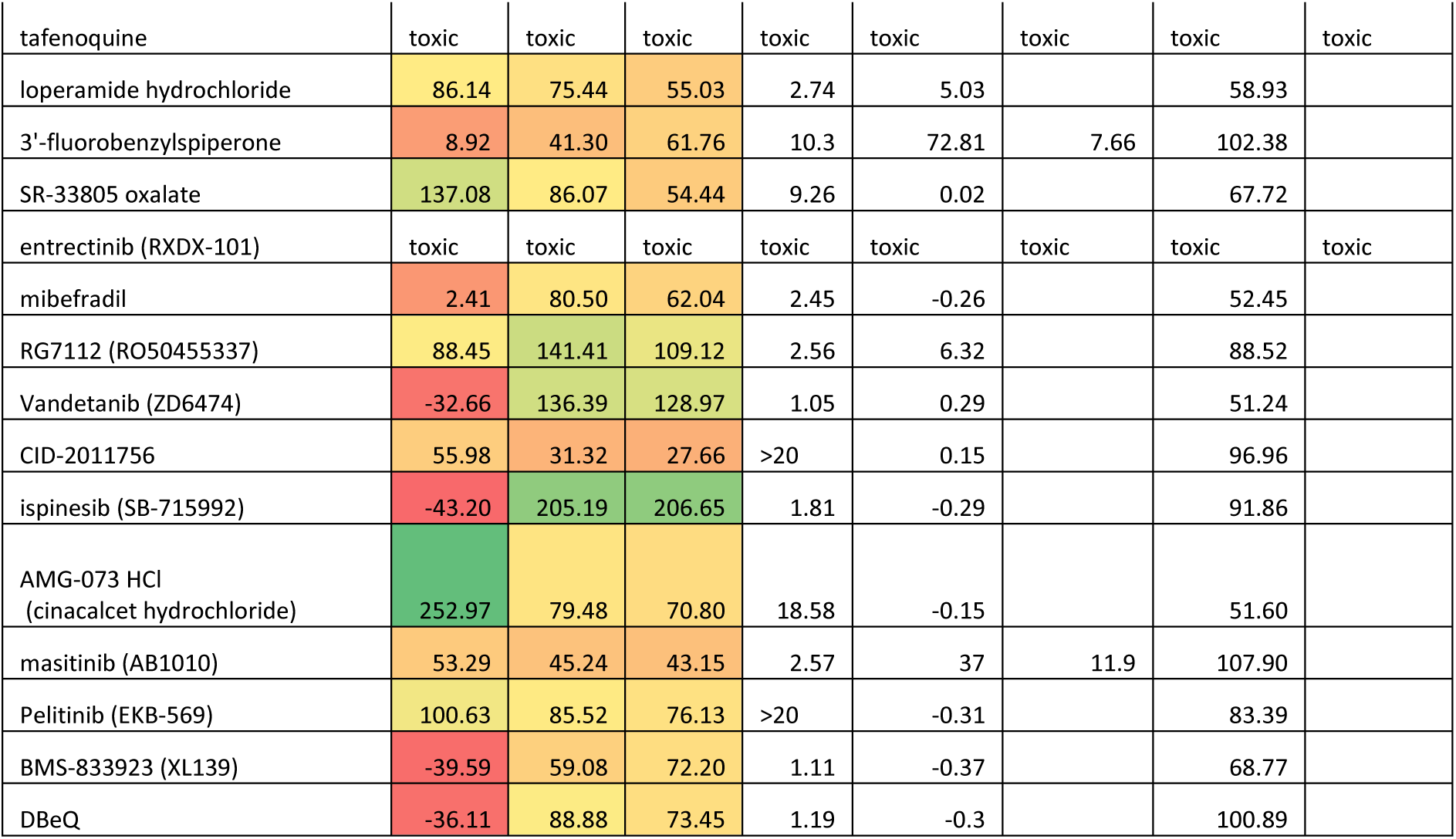
PLD EC_50_ determination of anti-SARS-CoV-2 CPE active compounds, correlation to available CPE EC_50_. PLD effect is normalized to amiodarone PLD induction effect at 10 µM, set to 100 %, DMSO is used as a negative control, set to 0 % PLD induction effect. Anti-SARS-CoV-2 CPE Data points are measured at 20 µM for infection in CaCo2 cells and at 10 µM for infection in Vero—E6 cells. Data points represent triplicate measurements. Empty fields: no data available

